# PPPred: Classifying Protein-phenotype Co-mentions Extracted from Biomedical Literature

**DOI:** 10.1101/654475

**Authors:** Morteza Pourreza Shahri, Mandi M. Roe, Gillian Reynolds, Indika Kahanda

## Abstract

The MEDLINE database provides an extensive source of scientific articles and heterogeneous biomedical information in the form of unstructured text. One of the most important knowledge present within articles are the relations between human proteins and their phenotypes, which can stay hidden due to the exponential growth of publications. This has presented a range of opportunities for the development of computational methods to extract these biomedical relations from the articles. However, currently, no such method exists for the automated extraction of relations involving human proteins and human phenotype ontology (HPO) terms. In our previous work, we developed a comprehensive database composed of all co-mentions of proteins and phenotypes. In this study, we present a supervised machine learning approach called PPPred (Protein-Phenotype Predictor) for classifying the validity of a given sentence-level co-mention. Using an in-house developed gold standard dataset, we demonstrate that PPPred significantly outperforms several baseline methods. This two-step approach of co-mention extraction and classification constitutes a complete biomedical relation extraction pipeline for extracting protein-phenotype relations.

**CCS CONCEPTS:** **•Computing methodologies** → **Information extraction; Supervised learning by classification;** •**Applied computing** →**Bioinformatics;**

## 1 INTRODUCTION

Proteins are one of the most critical biomolecules for the development and maintenance of life [3]. A cell’s full complement of expressed proteins, the proteome, is both dynamic and multidimensional with many proteins operating in a complex network ensuring the integrity of cellular structure and function [21]. Changes in critical regions of a protein’s structure often caused by errors in the underlying genetic sequence of the protein or in its regulation can alter the protein’s function-specific 3D structure, resulting in an alteration of phenotype [12]. In the medical context, a phenotype can be characterized as a deviation from normal morphology or behavior [34]. Well known alterations in phenotype brought about by changes in one or more proteins or their regulation involved in important biological pathways include Alzheimer’s disease, Parkinson’s disease, Huntington’s disease, cancer, cystic fibrosis and type II diabetes [3, 13, 26]. Uncovering novel changes in protein structure, function and regulation, and understanding how these alterations lead to human disorders is a very active area of research in the biological community [3, 5, 12, 13, 21, 26, 33, 37].

Human Phenotype Ontology (HPO) is a standardized vocabulary that includes a wide range of phenotypic abnormalities observed in human diseases [18]. HPO is composed of five sub-ontologies among which *Phenotypic abnormalities* is the main sub-ontology that describes clinical abnormalities. Each sub-ontology includes HPO terms and an associated HPO Identifiers (IDs), e.g. *Parkinsonism*, HP:0001300. Each sub-ontology is organized in a hierarchical structure where more general terms are close to the top while more specific terms are closer to the bottom. Each pair of terms in the hierarchy are linked with a *is-a* relationship. In this paper, we use *phenotypes* and *HPO terms*, interchangeably. HPO website^1^ provides gold-standard annotations for a large collection of human proteins through biocuration, which is the process of extracting knowledge from unstructured text and storing the data in knowledge bases. However, currently, only a small portion of known human proteins have HPO annotations [18]. But, it is believed that there are many other human proteins that are associated with diseases and hence should be annotated with HPO terms (Peter Robinson, personal communication, 2015).

Continuing to expand the knowledgebases such as HPO database through biocuration is of utmost importance for potential future downstream applications in medicine and healthcare. However, biocuration, which is usually performed manually with the help of computational tools [9], is generally considered tedious and resource-consuming. Hence, efficient and accurate computational tools are required to expedite the process in order to bridge the gap between the typically slower rate of human annotation versus the vast and exponentially-increasing amount of literature concerned with the subject [9]. As a result, developing computational models to extract relations between proteins and phenotypes has gained recent interest among scientists working in the field of biomedical natural language processing [11, 17, 19, 39]. However, to the best of our knowledge, no such computational methods exist for automatically extracting human protein-HPO term relations from biomedical literature.

As a solution to the above, we propose a two-step approach for extracting human protein-HPO term relations. The first step is to extract protein-HPO *co-mentions*, which are co-occurrences of protein names and phenotype names in a certain span of text i.e. a sentence, a paragraph, etc [17]. In our previous work, we developed ProPheno^2^, which is an online and publicly accessible dataset composed of proteins, phenotypes (HPO terms), and their co-occurrences (co-mentions) in text which are extracted from Medline abstracts and PubMed Central (PMC) Open Access full-text articles using a sophisticated in-house developed text mining pipeline [30]. This dataset covers all terms in the *Phenotypic abnormality* sub-ontology. However, a knowledge-free Natural Language Processing (NLP) pipeline extracts every co-mention of proteins and phenotypes, but not all protein-phenotype co-mentions simply imply that there is a relationship between the two entities (see Figure 1 for an example).

**Figure 1:**
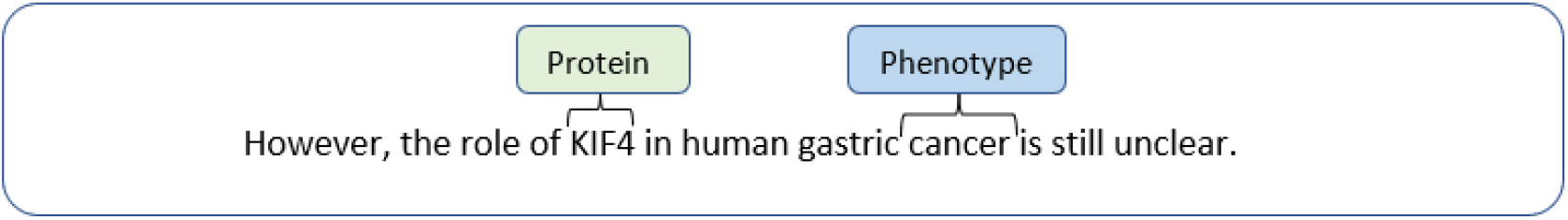
An example of a *bad* co-mention in which the sentence does not convey a relation between the protein, i.e. “KIF4”, and the phenotype, i.e. “cancer”. (PMID: 20711700)

Therefore, in the second step, extracted co-mentions are filtered using a co-mention classifier that can distinguish between good and bad co-mentions. We define a co-mention as a *good* co-mention if there is enough evidence conveyed in the corresponding span of text indicating a relationship between the protein and the phenotype. In other words, a good co-mention is a valid relationship between the two entities according to the meaning of the context text. Figure 2 depicts an example of a good co-mention of a protein and a phenotype in a sentence. The combination of a co-mention extractor and co-mention classifier/ filter constitutes a complete relation extraction pipeline.

**Figure 2:**
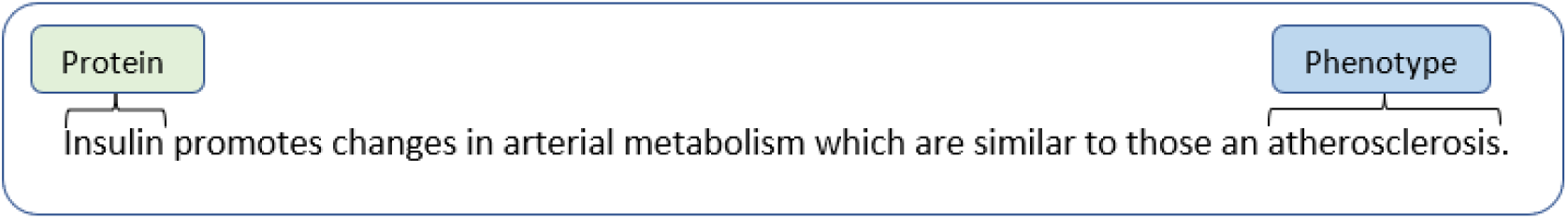
An example of a sentence-level protein-phenotype co-mention which is extracted from the article PMID: 798461.

The development of PPPred (Protein-Phenotype Predictor), a novel co-mention classifier for classifying protein-phenotype co-mentions, is the primary focus of the work presented in this paper. We first randomly select a subset of co-mentions from the ProPheno database and have them curated through two biologists. This gold-standard dataset is composed of 809 human protein-HPO term co-mentions annotated with binary labels of good/ bad. Then we use this gold-standard dataset for developing predictive models using machine learning techniques. Our machine learning models employ a large collection of both syntactic and semantic features. Finally, we demonstrate that PPPred significantly outperforms other baseline methods on the task of protein-HPO terms co-mention classification.

The main contributions of the paper are as follows. This is the first analysis of the problem of human protein-HPO term relation extraction from biomedical literature. We model this relation extraction task as a two-step process composed of co-mention extraction and classification. We formulate the co-mention classification problem as a supervised learning problem using the gold-standard data generated by biologists. This is also the first such gold-standard data for human protein-HPO term relation extraction and is made publicly available^3^. A filter or a classifier that could identify good co-mentions can be used by annotators to significantly speed up the biocuration process. In addition, this can be used to provide much higher quality co-mentions as input to other downstream applications such as human protein-HPO term prediction [29], which would likely lead to better predictions.

The rest of the paper is organized as follows. Section 2 provides a brief background on the related work in this area. The proposed method is discussed in Section 3. Section 4 discusses the results of running this method and compares the results with other methods and provides a discussion on the results. Finally, Section 5 concludes the study and discusses future work and open problems.

## 2 RELATED WORK

The main approaches for biomedical relation extraction include co-occurrence-based methods, rule-based methods, and machine learning-based methods. Co-occurrence methods simply look for any co-mention of the two entities of interest in a particular span of text, e.g. sentence, paragraph, etc., and usually provide low precision and high recall values [4]. Rule-based methods define linguistic patterns and extract the relations using the patterns [1, 23, 31]. The rules can be derived from manually annotated corpora using machine learning algorithms or defined manually by a domain expert. Machine learning-based approaches are also employed for the relation extraction from biomedical text [17, 20, 22, 38].

The machine learning category includes methods based on feature engineering, graph kernels, and deep learning. Support Vector Machines (SVMs) have shown high performance in biomedical relation extraction, but they need feature engineering which is a skill-dependent task [42]. Kernel-based methods also require designing suitable kernel functions. Deep neural network-based methods eliminate the need for feature extraction and defining rules, and provide state-of-the-art on various tasks in biomedical relation extraction [28, 42]. However, they typically require very large data sets compared to other traditional machine learning models.

Sekimizu et al. employ the most frequently seen verbs from Medline abstracts, and they try to find the subject and object terms for some of these verbs [36]. They linguistically analyze raw texts and then apply the mentioned method for classifying genes and gene products, and for identifying the relations between those entities. Temkin and Gilder propose a method based on using lexical analyzer and context-free grammar for extracting relations between genes, proteins, and small molecules from unstructured text [40]. Yakushiji et al. introduce a full parser for analyzing biomedical text using a general-purpose parser and grammar [41]. Coulet et al. also propose a system for extracting pharmacogenomics relations from biomedical text using a semantic network on relations [7].

Ng and Wong propose a prototype system based on pattern matching for automatic pathway discovery from abstracts [27]. They employ two sets of rules for specifying the patterns for protein name identification and for extracting protein-protein interactions (PPI). Huang et al. introduce a method based on dynamic programming for discovering patterns by aligning related sentences and key verbs, and they use this method to find protein-protein interactions from full-text articles [14].

Craven presents a machine learning method for mapping information from Medline abstracts to knowledge bases [8]. Katrenko and Adriaans propose a method that uses syntactic information and can be used with various machine learning methods [16]. Marcotte et al. introduce a Bayesian approach using the probability of discussing the PPI interactions using the frequency of 80 discriminating words from Medline abstracts [24]. Rosario and Hearst employ neural networks using lexical, syntactic, and semantic features for distinguishing seven relations between entities “disease” and “treatment” [35].

Rindflesch et al. present a natural language processing method for identifying casual relations between diseases and genetic phenomena [32]. Fundel et al. propose RelEx which is based on parse trees and simple rules and can be used for extracting relations between genes and proteins from Medline abstracts [10]. Bui et al. introduce an algorithm for extracting protein-protein interactions from biomedical literature based on the semantic properties of text and support vector machines for classifying PPI pairs [2].

Korbel et al. employ an unsupervised, systematic approach for finding relations between genes and phenotypic characteristics using Medline abstracts [19]. First, they retrieve abstracts that contain phenotypic similarities of species and then find genes that are present in the corresponding genomes. Goh et al. propose a method to find genotype-phenotype relations which combines molecular and phenotypic information [11]. Khordad and Mercer introduce a machine learning method for identifying genotype-phenotype relations which uses a semi-automatic approach for annotating more sentences to enlarge the training set [17].

Despite a large number of studies conducted on extracting entity relations from the biomedical literature (including a handful of methods for extracting relations between genes/proteins and phenotypes, no methods exist specifically for human protein-HPO term relation extraction. Therefore, to the best of our knowledge, this is the first study on the problem of protein-HPO term relation extraction from biomedical literature and the PPPred is the first such method. We note that GenePheno [15] is the only related method that uses an ontology-based approach to extract gene-phenotype associations from the literature. It first recognizes all mentions of gene and HPO terms within sentences in the whole corpora and then uses a co-occurrence based metric for ranking those pairs. Highest ranked pairs are predicted as gene-phenotype associations. While GenePheno does not predict top-ranked relations (i.e. sentences), we still use it as one of the baseline methods due to the close proximity of the problem solved by their method and the task of protein-phenotype co-mention classification addressed by PPPred.

## 3 METHODOLOGY

## 3.1 Approach

In this work, we formulate the task of co-mention classification as a supervised learning problem as described below.

Given a context *C* = *w*_1_*w*_2_..*e*_1_..*w*_3_..*e*_2_..*w*_*n* −1_*w*_*n*_ composed of words *w*_*i*_ and the two entities *e*_1_ and *e*_2_, we define a mapping *f*_*R*_(·) as:

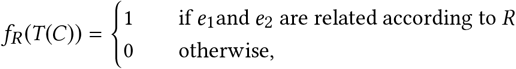

where *T (C)* is a high-level feature representation of the context, *e*_1_ and *e*_2_ are the entities representing the protein and the phenotype and *R* is the relation that represents the protein-phenotype relationship between the two. An example is considered a positive example if the meaning of the context suggests that the protein mentioned has this function (i.e. a good co-mention). Otherwise, it is labeled as a negative example.

In this work, the context *C* is a single sentence (i.e., the sentence containing the mentions of the two entities). Figure 2 depicts a sentence which is labeled as a positive example (i.e., *f*_*R*_ = 1) because it provides evidence for the relationship between the two entities “Insulin” (protein) and “Atherosclerosis” (phenotype). We model this problem as a supervised learning problem and use binary classifiers for learning *f*_*R*_.

Figure 3 depicts the overview of the PPPred pipeline, which is capable of classifying sentence-level co-mentions of proteins and phenotypes from biomedical literature. In this figure, we start by inputting a set of sentences that contain co-mentions of proteins and phenotypes. The preprocessing step is comprised of tokenization, removing punctuations and stop words, and stemming. In the next step, we extract features from the input sentences and train the model which is able to extract the relations. The steps are discussed in detail in the following sections.

**Figure 3:**
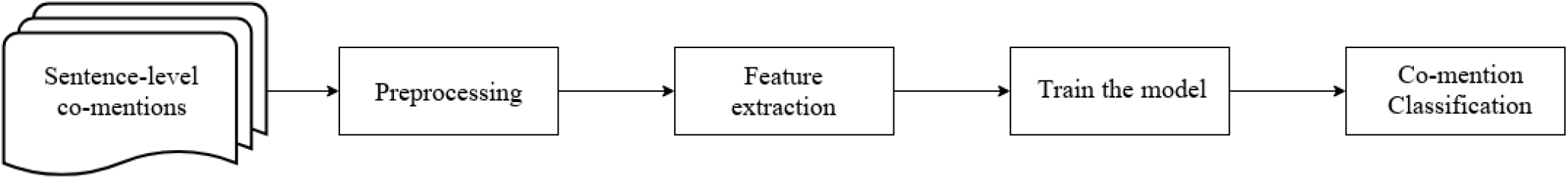
The Pipeline of the Proposed Method.

### 3.2 Dataset

The first step in building a co-mention classifier is to create a manually-annotated gold-standard dataset of co-mentions of proteins and phenotypes. For this purpose, we use ProPheno 1.0 [30], which is a dataset of proteins-phenotypes extracted from the entire biomedical literature. This dataset maps the proteins and phenotypes to the corresponding UniProt^4^ IDs and HPO IDs. We randomly select a dataset of 809 sentence-level co-mentions of proteins and phenotypes from ProPheno. This dataset is then annotated by two biologists to generate the gold-standard dataset. The annotators were provided instructions to label a co-mention as good/ positive if the sentence conveys that the protein and the phenotype has a relationship. Otherwise, the co-mention was labeled bad/negative.

Table 1 shows the distribution of co-mention types in the gold-standard dataset. According to the Table 1, 39% of sentences are extracted from the abstracts and 61% are from the full-text articles. Among the sentences from the abstracts, 53% are labeled as “good” and 47% are labeled as “bad”. The distribution for the sentences from the full-text articles is 70% and 30% “good” vs. “bad”, respectively. The overall class distribution is 64% and 36% for “good” and “bad”, respectively. The inter-annotator agreement is calculated using the Cohen’s Kappa statistic [25] and the corresponding value is 0.64 that shows substantial agreement.

**Table 1:** The class distribution in the gold standard dataset.

|  | Good | Bad | Total |
| --- | --- | --- | --- |
| Sentences from abstracts | 169 | 147 | 316 |
| Sentences from full-texts | 348 | 145 | 493 |
| All sentences | 517 | 292 | 809 |

Tables 2 and 3 show the most frequent phenotypes and proteins in the dataset, respectively. According to the tables, 15% of the sentences mention the protein “Receptor tyrosine-protein kinase erbB-2” (P04626) and 43% of the sentences discuss the HPO term “Neoplasm” (HP:0002664) (other names: “Cancer” or “Tumour”). Table 4 also demonstrates the most frequent protein-phenotype pairs mentioned in the dataset. We observe that 10% of the co-mentions in the dataset mention above protein-phenotype pair, which shows this pair is a well-studied protein-phenotype pair. Figure 4 depicts the distribution of the depths of HPO terms in the gold-standard.

**Table 2:** Most frequent HPO terms mentioned in the dataset.

| Index | HPO ID | HPO Term | Depth | Frequency |
| --- | --- | --- | --- | --- |
| 1 | HP:0002664 | Neoplasm | 2 | 348 |
| 2 | HP:0003002 | Breast carcinoma | 5 | 69 |
| 3 | HP:0001909 | Leukemia | 4 | 24 |
| 4 | HP:0002861 | Melanoma | 4 | 21 |
| 5 | HP:0000855 | Insulin resistance | 5 | 15 |

**Table 3:** Most frequent proteins mentioned in the dataset.

| Index | UniProt ID | Protein Name | Freq. |
| --- | --- | --- | --- |
| 1 | P04626 | Receptor tyrosine-protein kinase erbB-2 | 120 |
| 2 | Q9Y617 | Phosphoserine aminotransferase | 45 |
| 3 | O14788 | Tumor necrosis factor ligand superfamily member 11 | 23 |
| 4 | P01308 | Insulin | 20 |
| 5 | P03971 | Muellerian-inhibiting factor | 14 |

**Table 4:** Most frequent protein-HPO term pairs mentioned in the dataset.

| Index | UniProt ID | HPO ID | Protein Name | HPO Term | Frequency |
| --- | --- | --- | --- | --- | --- |
| 1 | P04626 | HP:0002664 | Receptor tyrosine-protein kinase erbB-2 | Neoplasm | 79 |
| 2 | Q9Y617 | HP:0002664 | Phosphoserine aminotransferase | Neoplasm | 42 |
| 3 | P04626 | HP:0003002 | Receptor tyrosine-protein kinase erbB-2 | Breast carcinoma | 31 |
| 4 | P09486 | HP:0002664 | SPARC | Neoplasm | 9 |
| 5 | P21860 | HP:0002664 | Receptor tyrosine-protein kinase erbB-3 | Neoplasm | 8 |

**Figure 4:**
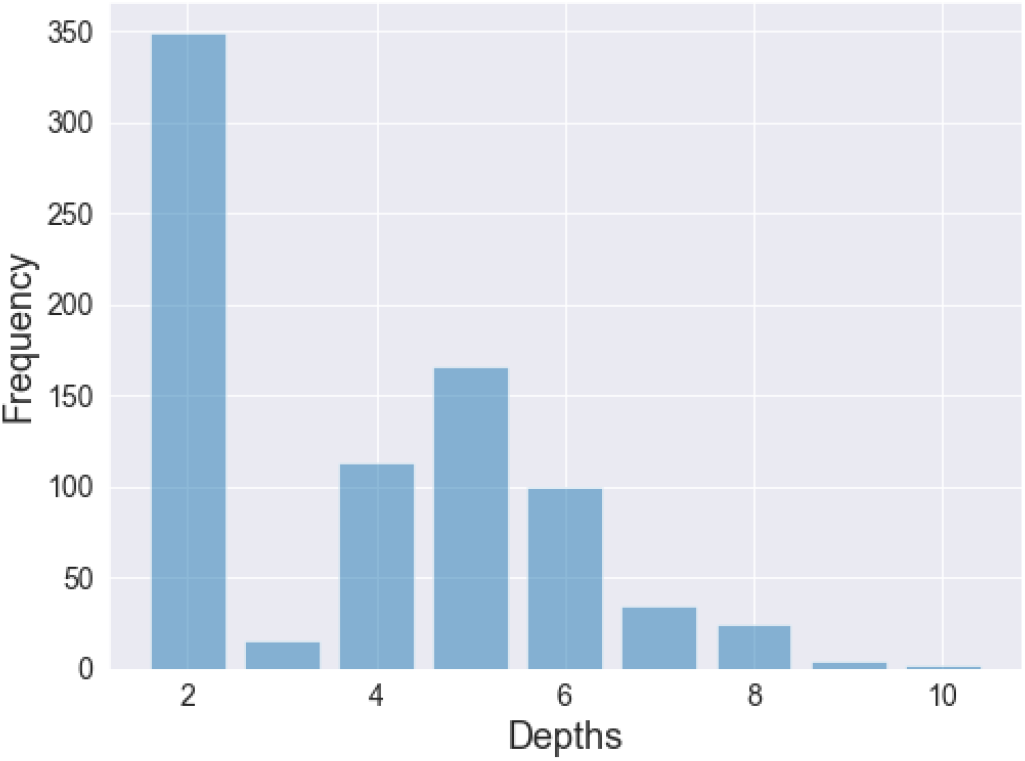
Distribution of depth of HPO terms in the annotated co-mention data.

### 3.3 Preprocessing

In the next step, we perform preprocessing on the sentences, which is basically employing tokenization, and removing highly frequent words from sentences (stop words), and also performing lemmatization. In this step, we replace protein and phenotype entities by PROT and PHENO, respectively. This replacement helps us to keep track of the actual labels when the sentence contains more than one entity with the same name and helps to avoid confusion when the entity names contain more than one word.

### 3.4 Feature Extraction

We define the following items as the features for classification. These features are categorized into three major types, i.e. bag-of-words, engineered features, and distantly supervised (DS) features.

#### 3.4.1 Bag-of-words (BoW) Feature

Here each feature is a token from the context sentence while the feature value is their corresponding frequency.

#### 3.4.2 Engineered Features

We obtained these features based on (1) domain expertise and (2) informative features used with similar relation extraction problem [22]. The full list of engineered features and their value type (within parentheses) is as follows:

1. Shortest dependency path between PROT and PHENO in the dependency graph of the sentence (integer).
2. The head words of PROT and PHENO in the sentences (string).
3. Part-of-speech tags of the entities and next tokens of entities in the sentences (string).
4. The number of tokens in sentences (integer).
5. Existence of interaction words acquired from a study by Chowdhary et al [6] (boolean).
6. Existence of seven trigger words provided by biologists, e.g. “provide”, “improve”, “confer”, etc (boolean).
7. Position of PROT in the sentence (integer).
8. Position of PHENO in the sentence (integer).
9. Tokens before and after PROT and PHENO (string).
10. Whether PROT is mentioned before PHENO in the sentence (boolean).
11. Existence of doubt in the sentence, e.g. “may”, “might”, etc (boolean).
12. Existence of negation words such as “no”, “not”, etc (boolean).

### 3.4.3 DS Features

We obtained the DS features by utilizing (1) the full set of co-mentions (i.e. unlabeled) available in ProPheno, and (2) the annotations available in the HPO database, which we call the silver-standard (SS). These features are listed in detail as follows:

1. Number of co-mentions containing the phenotype name (integer).
2. Number of co-mentions containing the protein name (integer).
3. Number of co-mentions containing both protein and phenotype name (integer).
4. Normalized number of co-mentions containing the protein name (float).
5. Normalized number of co-mentions containing the phenotype name (float).
6. Normalized number of co-mentions containing both protein and phenotype name (float).
7. Number of pair-specific co-mentions containing the protein name (integer).
8. Number of pair-specific co-mentions containing the phenotype name (integer).
9. Number of pair-specific co-mentions containing both protein and phenotype name (integer).
10. Normalized number of pair-specific co-mentions containing the protein name (float).
11. Normalized number of pair-specific co-mentions containing the phenotype name (float).
12. Normalized number of pair-specific co-mentions containing both protein and phenotype name (float).
13. Number of annotations in SS for the protein (integer).
14. Number of annotations in SS for the phenotype (integer).
15. Annotation score for the protein and the phenotype in SS (0 or 1).
16. Number of propagated annotations in SS for the protein (integer).
17. Number of propagated annotations in SS for the phenotype (integer).
18. Propagated annotation score for the protein and the phenotype in SS (0 or 1).

We normalize the number of co-mentions containing protein name, phenotype name, or a pair of protein-phenotype by dividing their frequencies by the number of unique articles that contain that specific protein, phenotype, or pair, respectively. We also propagate the HPO annotations upward toward the root nodes by using the *true path rule* that means if an HPO term has an annotation with a specific protein, all of its ancestors are also annotated with that protein.

### 3.5 Experimental Setup

The scikit-learn^5^ package is used for implementing the classifier functionality. We normalize the feature vectors using the *L2 norm*. In a preliminary analysis, we compared various supervised learning algorithms such as SVM, Naïve Bayes, Decision Trees, K-Nearest Neighbors (KNN), and Gradient Boosting Trees (GBT) using their default parameter settings. We select SVM with Linear kernel for the rest of our experiments. We perform 10-times 5-fold cross-validation The performances are reported primarily using F-max (the optimal F-1 value). Precision and Recall at F-max are presented as well.

We compare PPPred with three baselines: (1) a strict rule-based method (rule-based 1), (2) a lenient rule-based method (rule-based 2), and (3) GenePheno [15]. The rule-based 1 method was developed in-house by a biologist using broad domain knowledge of the language used when describing alterations in protein sequence, activity, regulation and the resulting phenotypic changes. Commonly used words for sequence-based alterations included “mutation”, “deletion” and “insertion”. For protein expression changes, the phrases “upregulation”, “upregulates”, “downregulation”, “down-regulates”, “over-expression”, “under-expression”, “switches off”, “switches on”, “amplifies” and “enhances” were chosen. For direct protein-phenotype relationship descriptions, the phrases “associated with”, “triggered by” and “caused by” were used. This method assigns a score of 1 to co-mentions satisfying at least one of the following rules (and 0 otherwise):

- PROT (upregulation/ downregulation/ over-expression/ under-expression/ mutation) causes/ does not cause/ is (not) associated with PHENO
- some other entity (upregulates/ downregulates/ silences/ inhibits/ switches off/ switches on/ triggers/ activates/ amplifies/ over-expresses/ under-expresses / enhances) PROT causing/ which causes/ which is associated with PHENO
- PEHNO is (not) associated with/ triggered/ caused by (up-regulation/ downregulation/ mutation/ deletion/ insertion/) in PROT
- Mutation/ deletion/ insertion in PROT causes/ is associated with PHENO

The rule-based 2 method is lenient than the rule-based 1 method because it only checks whether any of the keywords in the rule-based 1 method is in the sentences. In other words, this method assigns a score to a co-mention based on the keyword(s) present in the sentence. The order or the position of the keywords (with respect to PROT and PHENO entities) are not considered.

GenePheno [15] is an ontology-based text mining method for predicting gene-phenotype associations using literature. While acknowledging this is not an apples-to-apples comparison, we perform the following in order to adapt it as a baseline. For each co-mention in our gold-standard, if the corresponding pair of the protein and the phenotype exists in the pre-generated GenePheno output file^6^, we consider it as a positive prediction (otherwise negative). We incorporate the NPMI (Normalized Pointwise Mutual Information) scores provided by GenePheno for each co-mention as the confidence scores for the predictions. Note that due to the possibility of the GenePheno method having access to some or all of the co-mentions from our test set, the performance we report is likely an over-estimation.

## 4 RESULTS AND DISCUSSION

Table 5 demonstrates the F-max, precision at F-max, and recall at F-max values of various supervised learning algorithms. We observe that the Linear SVM and Gradient Boosting Trees algorithms achieve best the F-max value (0.8). In addition, the Decision Trees, Naïve Bayes, and K-Nearest Neighbors algorithms provide F-max values of 0.78, 0.79, and 0.78, respectively. However, by comparing the precision values, we realized that Linear SVM and Gradient Boosting Trees provide higher precision values. Since Linear SVM is one of the top models among all the models we compared, we use that for the rest of our experiments.

**Table 5:** Comparison of various machine learning algorithms using default parameter settings. Performance evaluated using 5-fold cross-validation and the results reported using F-max and precision/ recall at F-max.

| Model | Precision | Recall | F-max |
| --- | --- | --- | --- |
| Linear SVM | 0.69 | 0.95 | 0.8 |
| Decision Trees | 0.64 | 0.99 | 0.78 |
| Naïve Bayes | 0.67 | 0.97 | 0.79 |
| K-Nearest Neighbors | 0.64 | 1.0 | 0.78 |
| Gradient Boosting Trees | 0.69 | 0.95 | 0.8 |

Table 6 shows the comparison of the results of running PPPred against two rule-based methods and GenePheno. We observe that rule-based 2 and GenePheno obtain similar values for precision, recall, and F-max, whereas Linear SVM produces a higher F-max value. Linear SVM also achieves higher precision value than the rule-based 2 and GenePheno methods. Due to the lack of confidence scores for the rule-based 1 method, we report the F1-score instead of F-max. We performed the paired T-test on the values to compare the significance of the difference between F-max values. We observed that Linear SVM significantly outperforms other methods by achieving a p-value of 4.3E-13.

**Table 6:** Comparison of PPPred (uses SVMs with Linear kernel) against several baseline methods. Performance evaluated using 5-fold nested cross-validation and the results reported using F-max and precision/ recall at F-max. *F1 score reported in-place of F-max due to the lack of confidence scores for Rule-based 1 method.

| Method | Precision | Recall | F-max |
| --- | --- | --- | --- |
| Rule-based 1 | 0.71 | 0.26 | 0.38* |
| Rule-based 2 | 0.63 | 1.0 | 0.78 |
| GenePheno | 0.63 | 1.0 | 0.78 |
| Linear SVM | 0.69 | 0.95 | 0.8 |

Figure 5 provides a comparison between the effectiveness of various features on the sentences from the abstracts, full-text articles, and all sentences. The results suggest that we obtain better performance using the co-mentions from the sentences extracted from the full-text articles in comparison with the sentences extracted from the abstracts. The precision values of co-mentions extracted from full-text articles are higher than the values obtained by the abstracts. In other words, the co-mentions extracted from full-text articles could be a valuable source of information for relation extraction. The next observation is that BoW features often provide good performance in terms of precision, recall, and F-max that indicates the BoW features are an essential feature for relation extraction. Engineered features provide higher precision in comparison with DS features, whereas the DS features achieve higher recall values. This observation suggests that these two sets of features can be used as complementary features for relation extraction.

**Figure 5:**
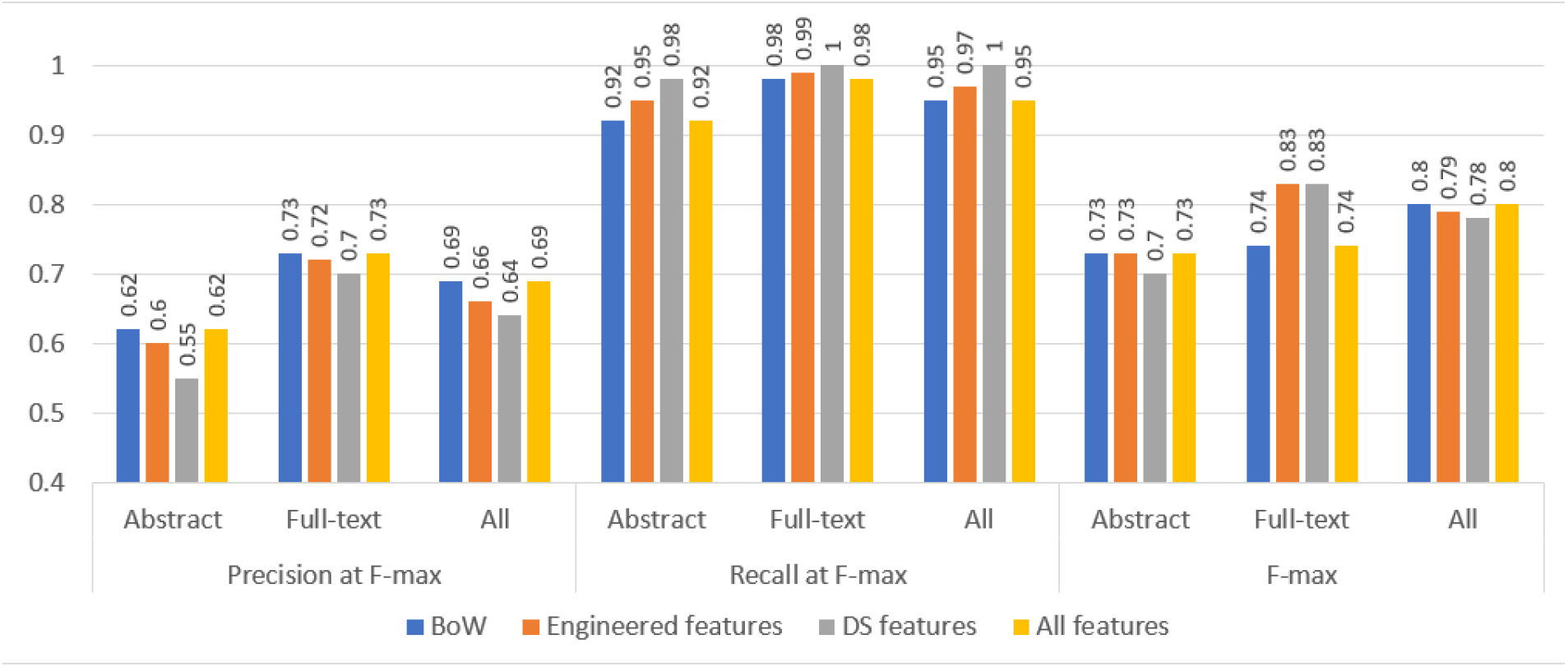
Comparison of the effectiveness of various features used with PPPred (uses SVMs with Linear kernel) on the sentences from abstracts, full-text articles, and all sentences. Performance evaluated using 5-fold nested cross-validation and the results reported using F-max and precision/ recall at F-max.

We investigate whether the training set suitably represents the problem by employing the learning curve with training sizes 20%-90% of the data and predicting on the holdout 10% of the data. Figure 6 depicts the learning curve with the mentioned training sizes. The increasing value of F-max shows that the dataset is under-representative of the problem and we need more training data.

**Figure 6:**
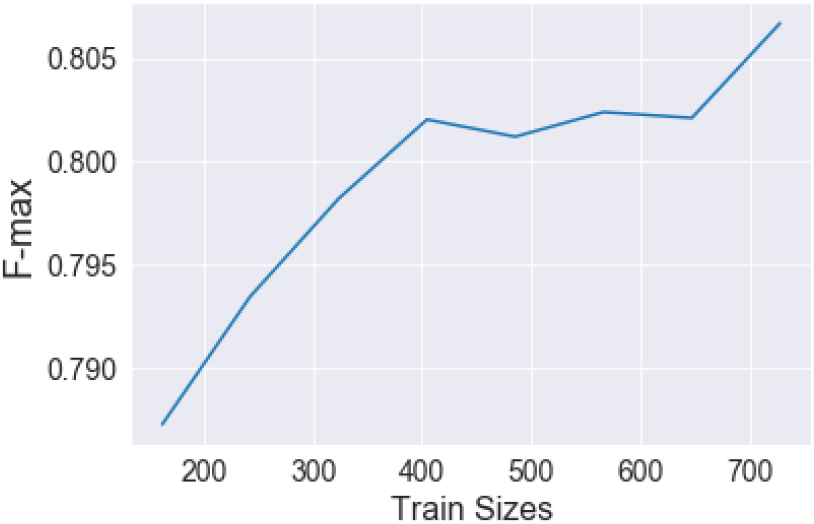
Learning Curve.

Relatively Low precision values using the Linear SVM algorithm suggest that we have many false positives. Therefore, to investigate the reason, we picked the top 5 false positives (sentences which are predicted as good with the highest confidence scores by the model whereas their actual labels are bad) shown in Table 7. We also picked the top 5 false negatives (co-mentions predicted as negatives with the lowest confidence scores, whereas their actual labels are good) which are shown in Table 8. By comparing sentences in Tables 8 and 7, we observe that the length of false negative and false positive sentences is similar and cannot be used as a criterion to differentiate between the co-mentions. Additionally, we observed that most of the phenotypes in the selected sentences are “cancer” or related to “cancer”. Therefore, the type of entities does not fully distinguish between good and bad co-mentions and requires further investigation.

**Table 7:**
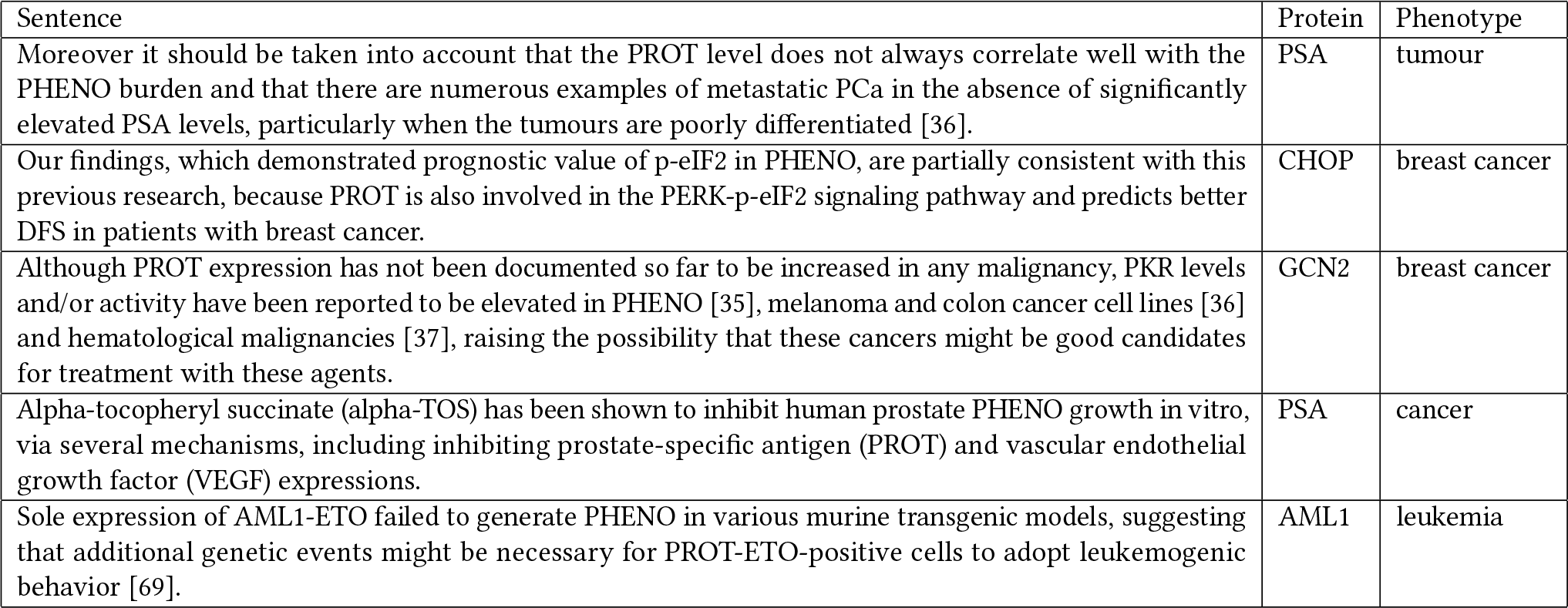
False Positives.

**Table 8:** False Negatives.

| Sentence | Protein | Phenotype |
| --- | --- | --- |
| Pedigree analyses of five families in which a form of spinocerebellar PHENO (SCA1) is present have been used to obtain additional information on the location of PROT on chromosome 6. | SCA1 | ataxia |
| The ratio of free to total PROT may increase the specificity of single serum PSA evaluations without decreasing its sensitivity for the diagnosis of prostate PHENO. | PSA | cancer |
| The PHENO cells are usually positive for cytokeratin 7 (CK7), epithelial membrane antigen (EMA), Cam 5.2, PROT, and mucicarmine stain while S100, Melan A, and human melanoma black-45 (HMB-45) highlight the non-neoplastic dendritic cells. | HER2 | tumour |
| Statistically significant changes in PROT signaling pathway activity between the xenograft and 2D cultures were also observed in MCF7 HER2-negative, ER-positive breast PHENO cell line and in the MDA-MB-231 and Hs578T triple negative breast cancer cell lines (GEO Series accession number GSE47583 for MCF7 and GSE36953 for MDA-MB-231 and Hs578T cell lines). | HER2 | cancer |
| The proportion of cases overexpressing PROT by tumor subtype was 72% for esophageal adenocarcinoma, 69% for gastric cardia adenocarcinoma, 52% for non-cardia gastric adenocarcinoma, and 67% for esophageal PHENO. | P53 | squamous cell carcinoma |

## 5 CONCLUSION AND FUTURE WORKS

In this project, we created a co-mention classifier/filter which is capable of distinguishing between good and bad co-mentions of proteins and phenotypes in sentences. We created a pipeline in which we perform preprocessing on manually-annotated sentence-level co-mentions of proteins and phenotypes, and by training a model on a set of features extracted from the sentences, we are able to classify the sentences comprising co-mentions of proteins and phenotypes. This classifier can be employed to perform relation extraction on protein and phenotype entities mentioned in biomedical literature. We observed that Linear SVM provides the best F-max score using five-fold cross-validation.

Nevertheless, there is still a lot of avenues to work in this area. We utilized syntactic features extracted from sentences, however, a potential future work is to use more specific syntactic features from sentences, e.g. the shape of the dependency graph. We also plan to do the classification on positive relations, negative relations, and no relations between entities to be able to extract more specific relations from biomedical literature by converting the problem into a multi-class classification. We also plan to apply deep learning and word embeddings to this dataset. We plan to incorporate the section titles, e.g. Introduction, Conclusion, etc., to employ only the more informative sentences. We also plan to utilize features based on the soft similarity between sentences and in the future, we are going to expand the study and include larger spans of text, i.e. paragraphs and documents.

## Footnotes

1 https://hpo.jax.org/app

2 http://propheno.cs.montana.edu

3 https://github.com/MSU-KAHANDA-LAB/protein-phenotype-relation-extraction

4 https://www.uniprot.org

5 https://scikit-learn.org/

6 https://zenodo.org/record/2532614

## REFERENCES

[1] Behrouz Bokharaeian, Alberto Diaz, Nasrin Taghizadeh, Hamidreza Chitsaz, and Ramyar Chavoshinejad. 2017. SNPPhenA: a corpus for extracting ranked associations of single-nucleotide polymorphisms and phenotypes from literature. Journal of biomedical semantics 8, 1 (2017), 14.

[2] Qyoc-Chinh Bui, Sophia Katrenko, and Peter MA Sloot. 2010. A hybrid approach to extract Protein–Protein Interactions. Bioinformatics 27, 2 (2010), 259–265.

[3] Sumit Kumar Chaturvedi, Mohammad Khursheed Siddiqi, Parvez Alam, and Rizwan Hasan Khan. 2016. Protein misfolding and aggregation: mechanism, factors and detection. Process Biochemistry 51, 9 (2016), 1183–1192.

[4] Elizabeth S Chen, George Hripcsak, Hua Xu, Marianthi Markatou, and Carol Friedman. 2008. Automated acquisition of disease–drug knowledge from biomedical and clinical documents: an initial study. Journal of the American Medical Informatics Association 15, 1 (2008), 87–98.

[5] Fabrizio Chiti and Christopher M Dobson. 2017. Protein misfolding, amyloid formation, and human disease: a summary of progress over the last decade. Annual review of biochemistry 86 (2017), 27–68.

[6] Rajesh Chowdhary, Jinfeng Zhang, and Jun S Liu. 2009. Bayesian inference of protein–protein interactions from biological literature. Bioinformatics 25, 12 (2009), 1536–1542.

[7] Adrien Coulet, Nigam H Shah, and others. 2010. Using text to build semantic networks for pharmacogenomics. Journal of biomedical informatics 43, 6 (2010), 1009–1019.

[8] Mark Craven. 1999. Learning to extract relations from MEDLINE. In AAAI-99 workshop on machine learning for information extraction, Vol. 5. The AAAI Press, 604–611.

[9] International Society for Biocuration. 2018. Biocuration: Distilling data into knowledge. PLOS Biology 16, 4 (04 2018), 1–8. DOI:http://dx.doi.org/10.1371/journal.pbio.2002846

[10] Katrin Fundel, Robert Kuffner, and Ralf Zimmer. 2006. RelExfi?Relation extraction using dependency parse trees. Bioinformatics 23, 3 (2006), 365–371.

[11] Chern-Sing Goh, Tara A Gianoulis, and others. 2006. Integration of curated databases to identify genotype-phenotype associations. BMC genomics 7, 1 (2006), 257.

[12] Peter W Harrison, Alison E Wright, and Judith E Mank. 2012. The evolution of gene expression and the transcriptome–phenotype relationship. In Seminars in cell & developmental biology, Vol. 23. Elsevier, 222–229.

[13] F Ulrich Hartl. 2017. Protein misfolding diseases. Annual Review of Biochemistry 86 (2017), 21–26.

[14] Minlie Huang, Xiaoyan Zhu, Yu Hao, Donald G Payan, Kunbin Qy, and Ming Li. 2004. Discovering patterns to extract Protein–Protein Interactions from full texts. Bioinformatics 20, 18 (2004), 3604–3612.

[15] Şenay Kafkas and Robert Hoehndorf. 2019. Ontology based text mining of gene-phenotype associations: application to candidate gene prediction. Database 2019 (2019).

[16] Sophia Katrenko and Pieter Adriaans. 2007. Learning relations from biomedical corpora using dependency trees. In Knowledge Discovery and Emergent Complexity in Bioinformatics. Springer, 61–80.

[17] Maryam Khordad and Robert E Mercer. 2017. Identifying genotype-phenotype relationships in biomedical text. Journal of biomedical semantics 8, 1 (2017), 57.

[18] Sebastian Köhler, Sandra C Doelken, and others. 2013. The Human Phenotype Ontology project: linking molecular biology and disease through phenotype data. Nucleic acids research 42, D1 (2013), D966–D974.

[19] Jan O Korbel, Tobias Doerks, and others. 2005. Systematic association of genes to phenotypes by genome and literature mining. PLoS biology 3, 5 (2005), e134.

[20] Andre Lamurias, Luka A Clarke, and Francisco M Couto. 2017. Extracting microRNA-gene relations from biomedical literature using distant supervision. PloS one 12, 3 (2017), e0171929.

[21] Mark Larance and Angus I Lamond. 2015. Multidimensional proteomics for cell biology. Nature reviews Molecular cell biology 16, 5 (2015), 269.

[22] Pei-Yau Lung, Zhe He, Tingting Zhao, Disa Yu, and Jinfeng Zhang. 2019. Extracting chemical–protein interactions from literature using sentence structure analysis and feature engineering. Database 2019 (2019).

[23] ASM Ashique Mahmood, Tsung-Jung Wu, Raja Mazumder, and K Vijay-Shanker. 2016. DiMeX: a text mining system for mutation-disease association extraction. PloS one 11, 4 (2016), e0152725.

[24] Edward M Marcotte, Ioannis Xenarios, and David Eisenberg. 2001. Mining literature for Protein–Protein Interactions. Bioinformatics 17, 4 (2001), 359–363.

[25] Mary L McHugh. 2012. Interrater reliability: the Kappa statistic. Biochemia medica: Biochemia medica 22, 3 (2012), 276–282.

[26] Ines Moreno-Gonzalez, George Edwards III, Natalia Salvadores, Mohammad Shahnawaz, Rodrigo Diaz-Espinoza, and Claudio Soto. 2017. Molecular interaction between type 2 diabetes and Alzheimerfis disease through cross-seeding of protein misfolding. Molecular psychiatry 22, 9 (2017), 1327.

[27] See-Kiong Ng and Marie Wong. 1999. Toward routine automatic pathway discovery from on-line scientific text abstracts. Genome Informatics 10 (1999), 104–112.

[28] Yifan Peng, Anthony Rios, Ramakanth Kavuluru, and Zhiyong Lu. 2018. Extracting chemical–protein relations with ensembles of SVM and deep learning models. Database 2018 (2018), bay073.

[29] Morteza Pourreza Shahri and Indika Kahanda. 2018. Extracting Co-mention Features from Biomedical Literature for Automated Protein Phenotype Prediction using PHENOstruct. In 10th International Conference on Bioinformatics and Computational Biology, BICOB 2018. 123–128.

[30] Morteza Pourreza Shahri and Indika Kahanda. 2019. ProPheno 1.0: An online dataset for accelerating the complete characterization of the human protein-phenotype landscape in biomedical literature. (2019). DOI:http://dx.doi.org/10.7287/peerj.preprints.27479v2

[31] KE Ravikumar, Majid Rastegar-Mojarad, and Hongfang Liu. 2017. BELMiner: adapting a rule-based relation extraction system to extract biological expression language statements from bio-medical literature evidence sentences. Database 2017 (2017).

[32] Thomas C Rindflesch, Bisharah Libbus, and others. 2003. Semantic relations asserting the etiology of genetic diseases. In *AMIA Annual Symposium Proceedings*, Vol. 2003. American Medical Informatics Association, 554.

[33] Marylyn D Ritchie, Emily R Holzinger, Ruowang Li, Sarah A Pendergrass, and Dokyoon Kim. 2015. Methods of integrating data to uncover genotype– phenotype interactions. Nature Reviews Genetics 16, 2 (2015), 85.

[34] Peter N Robinson. 2012. Deep phenotyping for precision medicine. Human mutation 33, 5 (2012), 777–780.

[35] Barbara Rosario and Marti A Hearst. 2004. Classifying semantic relations in bioscience texts. In Proceedings of the 42nd annual meeting on association for computational linguistics. Association for Computational Linguistics, 430.

[36] Takeshi Sekimizu, Hyun S Park, and Jun’ichi Tsujii. 1998. Identifying the interaction between genes and gene products based on frequently seen verbs in medline abstracts. Genome informatics 9 (1998), 62–71.

[37] Qiancheng Shen, Feixiong Cheng, Huili Song, Weiqiang Lu, Junfei Zhao, Xiaoli An, Mingyao Liu, Guoqiang Chen, Zhongming Zhao, and Jian Zhang. 2017. Proteome-scale investigation of protein allosteric regulation perturbed by somatic mutations in 7,000 cancer genomes. The American Journal of Human Genetics 100, 1 (2017), 5–20.

[38] Ayush Singhal, Michael Simmons, and Zhiyong Lu. 2016. Text mining for precision medicine: automating disease-mutation relationship extraction from biomedical literature. Journal of the American Medical Informatics Association 23, 4 (2016), 766–772.

[39] Ayush Singhal, Michael Simmons, and Zhiyong Lu. 2016. Text mining genotypephenotype relationships from biomedical literature for database curation and precision medicine. PLoS computational biology 12, 11 (2016), e1005017.

[40] Joshua M Temkin and Mark R Gilder. 2003. Extraction of protein interaction information from unstructured text using a context-free grammar. Bioinformatics 19, 16 (2003), 2046–2053.

[41] Akane Yakushiji, Yuka Tateisi, and others. 2000. Event extraction from biomedical papers using a full parser. In Biocomputing 2001. World Scientific, 408–419.

[42] Yijia Zhang, Hongfei Lin, Zhihao Yang, Jian Wang, Shaowu Zhang, Yuanyuan Sun, and Liang Yang. 2018. A hybrid model based on neural networks for biomedical relation extraction. Journal of biomedical informatics 81 (2018), 83–92.

